# Evolution of the latitudinal diversity gradient in the hyperdiverse ant genus *Pheidole*

**DOI:** 10.1101/295386

**Authors:** Evan P. Economo, Jen-Pan Huang, Georg Fischer, Eli M. Sarnat, Nitish Narula, Milan Janda, Benoit Guénard, John T. Longino, L. Lacey Knowles

**Affiliations:** Okinawa Institute of Science and Technology Graduate University, Onna, Okinawa, Japan, 904-0495; Department of Ecology & Evolutionary Biology, Museum of Zoology, University of Michigan, USA; National Laboratory for Ecological Analysis and Synthesis (LANASE), ENES, UNAM, Morelia, Mexico; Biology Centre of Czech Academy of Sciences, Ceske Budejovice, Czech Republic; The University of Hong Kong, School of Biological Sciences, Hong Kong, SAR, China.; Department of Biology, University of Utah, USA

**Keywords:** ants, latitudinal diversity gradient, tropical conservatism, diversification rate, diversity regulation, macroevolution

## Abstract

**Aim:** The latitudinal diversity gradient is the dominant pattern of life on Earth, but a consensus understanding of its origins has remained elusive. The analysis of recently diverged, hyper-rich invertebrate groups provides an opportunity to investigate latitudinal patterns with the statistical power of large trees while minimizing potentially confounding variation in ecology and history. Here, we synthesize global phylogenetic and macroecological data on a hyperdiverse (>1100 species) ant radiation, *Pheidole*, and evaluate the roles of three general explanations for the latitudinal gradient: variation in diversification rate, tropical conservatism, and ecological regulation.

**Location:** Global.

**Time Period:** The past 35 million years.

**Major taxa studied:** The hyperdiverse ant genus *Pheidole* Westwood.

**Methods:** We assembled geographic data for 1499 species and morphospecies, and inferred a dated phylogeny of *Pheidole* of 449 species, including 150 species newly sequenced for this study. We tested correlations between diversification rate and latitude with BAMM, HiSSE, GeoSSE, and FiSSE, and examined patterns of diversification as *Pheidole* spread around the globe.

**Results:** We found that *Pheidole* diversification occurred in series of bursts when new continents were colonized, followed by a slowdown in each region. There was no evidence of systematic variation of net diversification rates with latitude across any of the methods. Additionally, we found latitudinal affinity is moderately conserved with a Neotropical ancestor and phylogenetic inertia alone is sufficient to produce the gradient pattern.

**Main Conclusions:** Overall our results are consistent with tropical conservatism explaining the diversity gradient, while providing no evidence that diversification rate varies systematically with latitude. There is evidence of ecological regulation on continental scales through the pattern of diversification after colonization. These results shed light on the mechanisms underlying the diversity gradient, while contributing toward a much-needed invertebrate perspective on global biodiversity dynamics.

## INTRODUCTION

Understanding how ecological and evolutionary processes interact with historical factors to shape global biodiversity patterns remains a major goal of biology. The latitudinal diversity gradient (LDG) is the most general biogeographic pattern, yet we still lack a consensus understanding of its mechanisms (Pianka, 1966; Willig *et al.*, 2003; Mittelbach *et al.*, 2007; Fine, 2015). This is likely because many biological, physical, and historical factors that could plausibly affect diversity vary systematically with latitude, and thus a large number of hypotheses have been developed to explain the pattern. However, testing the predictions of different hypotheses empirically and evaluating their relative merits has proven to be a challenge.

Recently, the synthesis of large-scale geographic datasets along with large-scale phylogenetic data has provided new opportunities for empirical evaluation of hypotheses for the mechanisms underlying the LDG. These tests have mainly focused on vertebrates (e.g. Cardillo *et al.*, 2005; Weir & Schluter, 2007; Jetz *et al.*, 2012; Pyron & Wiens, 2013; Rolland *et al.*, 2014; Duchêne & Cardillo, 2015; Siqueira et al. 2016; Pigot *et al.*, 2016) and woody plants (Kerkhoff *et al.*, 2014), since those are the taxa with large-scale comprehensive data available. Several pioneering studies have examined latitudinal diversification patterns in insects (e.g. McKenna & Farrell, 2006; Hawkins & DeVries, 2009; Condamine *et al.*, 2012; Moreau & Bell, 2013; Pie, 2016; Owens *et al.*, 2017) although data-deficiency of most invertebrate groups makes taxonomic and/or geographic scope a problem for analysis.

Among invertebrates, ants are emerging as an exemplar taxon for global biodiversity studies. Ants are ecologically dominant in most terrestrial ecosystems and are, for an insect group, relatively well documented scientifically. Moreover, their diversity is high, but not intractably so, with richness on the same order as major vertebrate groups (~15,000 described ant species), and exhibit a marked latitudinal gradient (Kaspari *et al.*, 2004; Dunn *et al.*,2009). Recently, a new comprehensive dataset was compiled which gives the known geographic distribution of all described ant species across >400 geographic regions around the globe (Guénard *et al.*, 2017). These data, combined with progress toward reconstructing the ant tree of life (Brady *et al.*, 2006; Moreau *et al.*, 2006; Moreau & Bell, 2013; Ward *et al.*,2015), allow for inferences of the evolutionary underpinnings of large-scale diversity patterns in ants.

Here, we use globally distributed, hyperdiverse (>1100 described species) ant genus *Pheidole* as a model taxon to test hypotheses for the latitudinal diversity gradient. While over a hundred hypotheses have been proposed to explain the gradient (Willig *et al.*, 2003; Mittelbach *et al.*, 2007; Fine, 2015), these can roughly be sorted under three umbrella hypotheses which we use to frame our study: i) the *Diversification Rate hypothesis* (DRH), ii) the *Tropical Conservatism Hypothesis* (TCH), and iii) the *Ecological Regulation Hypothesis*(ERH).

First, the *Diversification Rate Hypothesis* posits that there is some causal factor that affects speciation and/or extinction rates and varies with latitude (e.g. reviewed in Pianka, 1966; Mittelbach *et al.*, 2007; Fine, 2015). This leads to a latitudinal disparity in species accumulation rate that underlies the gradient, rather than any regulation of total species numbers. Many such potential causal factors have been proposed. For example, temperature may affect mutation rates, which in turn could affect the rates of evolution of reproductive incompatibilities (Rohde, 1992). Such hypotheses about speciation contribute to the idea that faster origination in the tropics and dispersal outward lead to differential accumulation rates across latitude: the “out of the tropics” model (Jablonski *et al.*, 2006). Or, extinction rates could be higher in the temperate than tropical zone to due greater climatic variability (Weir & Schluter, 2007; Rolland *et al.*, 2014). The prediction of the DRH is straightforward: net diversification rate inferred from a phylogeny should be higher in tropical lineages compared with extratropical lineages.

Second, the *Tropical Conservatism Hypothesis* (TCH) (Pianka, 1966; Wiens & Donoghue, 2004) posits that the relative youth of colder temperate biomes combined with the inertia of phylogenetic niche conservatism (Wiens & Graham, 2005; Losos, 2008) has limited the accumulation of diversity in the temperate zone. In this scenario, net diversification rates do not necessarily vary with latitude, and the difference in richness is mainly due to time for diversification (Stephens & Wiens, 2003). This idea is based on the fact that historically the Earth has been much warmer than it is now, and much of what is now the temperate zone was covered by “megathermal” biomes. This hypothesis is supported by the fossil record; many lineages that used to occur in the Palearctic are now limited to tropical latitudes. This is true for ants as well; the Baltic amber ant fauna from the late Eocene has greater affinity to modern Indo-Australian faunas than modern Palaearctic faunas (Guénard *et al.*, 2015). The main prediction of this hypothesis is that the ancestral region of most groups is the tropics, transitions out of the tropical zone are rare, and thus the temperate clades are younger and nested within tropical clades. The fact that transition from the tropical to the temperate zones should be difficult because of the many nontrivial adaptations that ectothermal organisms such as ants need to overwinter in the higher latitudes. One additional key prediction of the TCH is that temperate diversification matches the timing of global cooling: specifically, that diversification of cold-adapted lineages accelerated after the Oligocene cooling 34mya.

Finally, the *Ecological Regulation Hypothesis* (ERH) posits that, due to some causal factor that varies with latitude, more species can coexist locally and regionally in tropical ecosystems than in temperate ecosystems. In this case, diversity is saturated at or near some ecological limit, and this “carrying capacity” of species varies with latitude regulating diversity from the top-down (Rabosky & Hurlbert, 2015). This is perhaps due to limitations on species coexistence that are driven by productivity or other factors (Pianka, 1966; Hurlbert & Stegen, 2014b). Speciation and extinction rates may vary over time to regulate richness at the requisite quota for a geographic region, but are not causally responsible for the disparity in diversity. Likewise, latitudinal affinity may be highly conserved or evolve quickly, but it would be immaterial to the origins of the gradient if diversity is regulated at levels that vary with latitude.

In a parallel study, Economo et al. (In Press) examined latitudinal patterns across 262 ant clades and tested hypotheses for the latitudinal gradient. That taxon-wide analysis focused on deeper timescales, and lacks phylogenetic resolution within recent radiations. They found that tropical lineages are more ancient than extratropical lineages, which mainly arose since the Oligocene cooling (past 34my), consistent with the TCH. Further, they found that diversification rate is highly heterogeneous but uncorrelated with latitude among ant clades, inconsistent with the DRH. Due to the limitations of phylogenetic data at such broad scales, they could not explicitly test for ecological regulation (ERH).

As with other studies on broad taxonomic scales (e.g. birds: Jetz *et al.*, 2012, mammals: Buckley & Jetz, 2007; Rolland *et al.*, 2014), the analyses across all ants provide the advantages of the statistical power of large numbers and a deep-time perspective. However, as many ecological, functional trait, and historical factors could contribute to variation in macroevolutionary rates at these deeper phylogenetic scales, this variation among clades may obscure underlying latitudinal effects that could be detectable among more similar, closely related lineages. For example, ant diversification has been shown to be related to functional traits (Blanchard & Moreau, 2017). Moreover, latitudinal gradients are often present within individual clades that evolved recently (Economo *et al.*, 2015a), and macrophylogenetic studies may miss the relevant scale of variation. Thus, the analysis of closely related lineages within younger, hyper-rich radiations provides a necessary complement to large-scale taxon-wide studies. In highly diverse groups, these radiations can provide both the statistical power of large numbers, while controlling to some degree for differences in ecology, functional traits, and historical factors.

The global radiation of *Pheidole* arose entirely since the Oligocene cooling (last 34my), during which time it has evolved a latitudinal gradient echoing the pattern for all ants (Economo *et al.*, 2015a). Thus, *Pheidole* presents an opportunity to examine diversification dynamics in this most recent period since the Oligocene, a period where many ant lineages transitioned out of the tropics, complementing the deeper timescales of the ant-wide study. While the low number of older extratropical ant lineages is consistent with the TCH (Economo, In Press), there is still an open question of whether niche conservatism or diversification rate differences explain the emergence of the gradient since the Oligocene. According to the TCH, the tropical ancestry of *Pheidole* combined with phylogenetic niche conservatism is sufficient to explain why there are more species in the tropics. The DRH predicts that *Pheidole* diversified more rapidly in the tropics, and this explains the disparity in diversity. Finally, we examine whether there is evidence for ecological limits to *Pheidole* diversification (ERH) by examining whether the radiation is undergoing pulse-like bursts of diversification. If so, we would expect a series of diversification pulses followed by slowdowns as the genus colonized different regions around the globe.

Here, we reconstruct a new global *Pheidole* phylogeny—the most comprehensive to date—increasing substantially the taxonomic and geographic coverage from previous studies of the genus (Moreau, 2008; Sarnat & Moreau, 2011; Economo *et al.*, 2015a; Economo *et al.*, 2015b). We use the new phylogeny and geographic data from the GABI database to test predictions of the three umbrella hypotheses for the latitudinal gradient. There is no biological reason why two or more of these mechanisms cannot be simultaneously operating (e.g. diversity can be regulated and speciation rate can vary systematically with latitude, or niche conservatism can occur along with diversity regulation). Thus, our goal is to rule out hypotheses rather than isolate a single exclusive answer. The analysis of this famously hyperdiverse radiation will advance our general understanding of the latitudinal gradient, the most pervasive pattern of life on Earth.

## METHODS

### Geographic Data

Our geographic data are based primarily on the Global Ant Biodiversity Informatics Project (GABI) database (Guénard *et al.*, 2017) which can be viewed through the website *antmaps.org* (Janicki *et al.*, 2016), and secondarily on the personal collection records of the authors. The former focuses on described species, while the second was used to supplement data on morphospecies for taxa included in the phylogenetic analysis.

### Phylogeny reconstruction

#### Taxon Selection

Compared with many other large ant radiations, the effort to reconstruct the phylogenetic history of *Pheidole* is relatively far along. A series of studies, beginning with Moreau 2008 (Moreau, 2008) and followed by other studies (Sarnat & Moreau, 2011; Economo *et al.*, 2015a; Economo *et al.*, 2015b) has produced a broad picture of the evolutionary history of the genus. However, for the purposes of understanding geographic patterns of diversification, having a larger, and more proportionally sampled phylogeny will provide additional statistical power and more robust results. Thus, we continued sampling *Pheidole* taxa for sequencing, focusing on sampling more taxa from the Neotropics, Madagascar, and SE Asia, which had been undersampled in previous analyses. In all, we increased the number of species from 282 taxa in the most recent global *Pheidole* phylogeny (Economo *et al.*, 2015a) to 449 taxa in the current contribution (Table S2).

#### Estimation of Sampling Completeness

One source of uncertainty in large-scale analyses of diversity is bias in taxonomic completeness overall and among different areas, particularly in relatively poorly known groups such as insects. While there is still a pronounced latitudinal gradient in *Pheidole* even among described species, there are undoubtedly many undescribed species in the genus, and it is probable they are disproportionately found in the tropics. While accounting for unobserved species is a challenge in any analysis, we devised an approximate method to calculate sampling completeness across areas given the information in hand, and use these estimates in our analysis of diversification rate. The details of our calculation are in the Supporting Information.

#### DNA Sequencing

Previous molecular work on *Pheidole* has generated a dataset based on eight nuclear loci [His3.3B (histone H3.3B F1 copy), Lop1 (long wavelength sensitive opsin 1), GRIK2 (glutamate receptor ionotropic, kainate 2-like), unc_4 (unc-4 homeodomain gene), LOC15 (uncharacterized locus LOC15), CAD (carbomoylphosphate synthase), EF-1α F2 (elongation factor 1-alpha F2), Top1 (DNA topoisomerase 1)], and one mitochondrial locus [CO1 (cytochrome oxidase 1)]. In a previous study (Economo et al. 2015a), all 9 loci were sequenced for a subset of 65 taxa representing the main *Pheidole* lineages around the world, while three loci (COI, Lop1, and His3.3B) were sequenced for all taxa to fill out the clades (217 taxa). This hierarchically redundant sampling design was chosen for reasons cost and time efficiency and to maximize the number of taxa, combined with the fact that many of the slow-evolving nuclear genes provide less information on recent divergences.

We added 167 new *Pheidole* taxa to this existing dataset by extending this sampling design and sequencing COI, Lop1, and His3.3B. We did not plan to sequence all 9 loci unless we found novel divergent clades not represented by taxa with all 9 genes sequenced in the earlier study (and we did not). Ant samples from field collections fixed in 95% EtOH were extracted for DNAs using DNeasy Blood & Tissue Kit (Qiagen, Hilden, Germany). The whole ant body was incubated in the extraction buffer without grinding during the first step, and then the complete ant specimen was removed before filtering and cleaning the extracts via a provided column. Extracted DNAs were subsequently used for PCR reactions for one mitochondrial (CO1) (Folmer *et al.*, 1994) and two nuclear (His3.3B and Lop1) regions. Each reaction contained 0.5 ul of extracted DNA, 1ul of 10 × buffer, 0.75 ul of MgCl2, 0.5 ul of 10mN dNTPs, 0.2 ul of 1% BSA, 0.4ul of of each primer, 0.04ul of Tag DNA polymerase (Invitrogen, USA), and ddH_2_O to make a total of 10 ul reaction. Standard PCR procedures were employed with annealing temperatures of 52, 60, and 60 C for CO1, His3.3B, and Lop1 regions, respectively. The amplicons were sequenced via a ABI^3700^ machine by the Sequencing Core at the University of Michigan. Sequences were checked using SeqMan (DNAStar Inc., USA).

#### Phylogenetic tree inference

We used Bayesian methods to infer a dated *Pheidole* phylogeny including 449 ingroup taxa (Table S2). To generate codon-aware alignments for these loci, we first searched NCBI’s non-redundant CDS database (Clark *et al.*, 2016) for reliable amino acid sequences for all loci and retrieved such sequences for seven of the nine loci with the following accession numbers: AIM2284.1 (CAD), ABW70333.1(CO1), EZA53539.1 (EF-1α F2), EGI60526.1 (His3.3B), ABW36758.1 (Lop1), EGI59282.1 (unc-4), and AIM43286.1 (Top1). These sequences were used as references for generating codon-aware alignments.

The CAD, unc-4, and Top1 alignments generated using MAFFT v7.205 (Katoh & Standley, 2013) (--retree 4; --maxiterate 1000) showed no frameshift mutations and/or insertions and deletions. However, the CO1, EF-1α F2, His3.3B, and Lop1 alignments did not match the reference sequences, showing disruptions in the translated amino acid alignments (such as the presence of numerous stop codons). For these loci, we used a codon-aware alignment software, MACSE v1.01b (Ranwez *et al.*, 2011), to generate the alignments. Reverse translations of the reliable amino acid reference sequences, accounting for all possibilities at each codon position, were passed as reliable input sequences to the software, we were able to assign codon positions within the exons in these seven loci. The resulting alignments were manually inspected and cleaned using Geneious R8 software. Furthermore, we identified, extracted, and separately aligned intronic regions wherever necessary. The remaining two loci, LOC15 and GRIK-2, were aligned using MAFFT. We concatenated all nine alignments and once again manually cleaned the master alignment, resulting in an alignment containing 8839 sites.

We used PartitionFinder v1.1.1 (Lanfear *et al.*, 2012) to determine the partitioning scheme and corresponding models of molecular evolution. The model scope included HKY, HKY+Γ, SYM, SYM+Γ, GTR, GTR+Γ, TrN, TrN+Γ, K80, K80+Γ, TrNef, TrNef+Γ, JC, and JC+Γ, branch lengths were set to ‘linked’, and model selection and comparison was set to Bayesian Information Criterion (BIC). PartitionFinder identified an optimal scheme containing 16 partitions (Table S3). We used ClockstaR (Duchene *et al.*, 2014) to determine the optimal number of clock models across our partitions for relaxed-clock phylogenetics, and a single linked clock was preferred based on the SEMmax criterion.

Our primary phylogenetic inference was conducted in BEAST2 v2.1.3 (Bouckaert *et al.*, 2014), but we first performed maximum likelihood (ML) reconstruction in RAxML v8.0.25 (Stamatakis, 2014). Using the partitioning scheme described above and the GTR+Γ model, we ran 75 ML inferences with 1000 bootstraps to find the ML tree. Using the *chronos* function in the *ape* package in R (Paradis *et al.*, 2004), we scaled the tree by calibrating the root node to a range of 50-60my. This tree was used as the starting tree for the BEAST2 analyses, but the topology was not fixed.

Unfortunately there are no reliable fossil calibrations available to date nodes *within* the genus. Thus, the age of the group can only be informed by the age of the stem node and information from fossils in related taxa across the subfamily Myrmicinae. Thus, because our analysis is concentrated within *Pheidole*, we preferred to use the stem node age distribution (i.e. the most recent common ancestor of *Pheidole* and its sister lineage *Cephalotes+Procryptocerus*) inferred as in a much larger analysis of the subfamily Myrmicinae (Ward *et al.*, 2015) that could make use of a broad range of molecular and fossil data to date the ages. Following those results, the stem node calibration was set to a normal distribution (mean: 58.0 mya, sigma, 4.8my) to match results from that study. Future work analyzing the global fossil record in *Pheidole* and placing fossil taxa within the *Pheidole* tree represents an important avenue for future phylogenetic work on the genus. Despite this limitation, the analyses in this paper depend mostly on relative, rather than absolute ages, and we draw no conclusions based on the precise timing of nodes in the tree.

We used a relaxed lognormal clock model linked across partitions (due to ClockstaR results), and used the partitioning scheme and models identified with PartitionFinder. Six independent analyses were run and chains were stopped between 45 and 80 million generations, after we observed convergence using Tracer software v1.6.0 (Rambaut 2014). We discarded the leading 33% of saved states as burnin, combined the remaining trees from all runs to create the posterior set, and generated the Maximum Clade Credibility tree and nodes set to median height. After pruning the outgroup, this tree was used for all subsequent analyses.

### Macroevolutionary rate inference

We took several complimentary approaches to estimating macroevolutionary rates and potential dependencies on latitude, primarily basing our anlaysis on BAMM (Bayesian Analysis of Macroevolutionary Mixtures, Rabosky, 2014), HiSSE (Hidden State Speciation and Extinction, Beaulieu & O’Meara, 2016), and FiSSE (Rabosky & Goldberg, 2017), with a secondary analysis using GeoSSE (Geographic State Speciation and Extinction, Goldberg *et al.*, 2011). These methods each have their strengths and weaknesses thus our approach is to use them collectively to seek conclusions about our data that are robust to methodological assumptions and implementation.

The main advantage of BAMM is that complex mixture models can be assessed with rate shifts across the tree, including accelerating and decelerating diversification rates. While trait-dependent diversification models are not fit directly, trait-diversification correlations can be assessed *post hoc* using structured rate permutations that estimate correlations while accounting for phylogenetic dependency (Rabosky & Huang, 2015). We use BAMM to test for correlations between latitude and net diversification rate, and evaluate evidence of decelerating diversification (ecological regulation of diversity) overall and in relation to the colonization of continents.

While BAMM has strengths in inferring complex mixtures of diversification processes, they are not explicitly trait-dependent, and the SSE family of methods explicitly fits models of trait-dependent diversification. SSE methods have been developed with different kinds of trait data, either based on binary traits (BISSE, Maddison *et al.*, 2007), continuous traits (QuaSSE, FitzJohn, 2010), or explicitly geographic traits (GeoSSE, Goldberg *et al.*, 2011). While these methods are explicitly for inferring trait-dependent speciation and extinction, they have the problem that differences in the focal trait are the only mechanisms that can cause shifts in macroevolutionary rates. If the real process has complex rate shifts then a more complex trait dependent model may fit better than a homogeneous null model, even if the shifts are not related to the traits per se, leading to type-I errors (Rabosky & Goldberg, 2015). These problems are at least partially solved by HiSSE (Beaulieu & O’Meara, 2016), a method that fits binary trait-dependent speciation and extinction models that can be formally tested against similarly complex trait-independent models. We thus primarily used HiSSE for our analysis. Since GeoSSE has been implemented for explicitly geographic dynamics, we also fit that model as a secondary test and present that analysis in the supplement.

Finally, as an additional test for variation in speciation rate with latitude, we used a non-parametric method, FiSSE (Rabosky & Goldberg, 2017), that does not depend on an assumed model structure and is robust to false inferences of trait-dependent evolution given a range of underlying complex evolutionary dynamics. FiSSE is limited to testing speciation rate differences it does not directly test for diversification rate differences. However, many (but not all) hypotheses for why diversification rate could vary with latitude are based on mechanisms acting on speciation rate (e.g. out-of-the-tropics model is one), so it is a partial test of the broader Diversification Rate Hypothesis.

#### BAMM implementation

We estimated net-diversification, speciation, and extinction rates through time for the inferred *Pheidole* tree using the program BAMM V2.5. The initial values for speciation rate, rate shift, and extinction rate were estimated using the setBAMMpriors function from the R package BAMMtools (Rabosky *et al.*, 2014) and specified in the BAMM control file. Specifically, a total of 2 × 10^8^ generations of rjMCMC searches with samples stored every 8000 generations were performed using speciation-extinction. A total of 1000 post burnin samples (50%) were retained. We performed two BAMM runs for each of three assumptions about sampling completeness (L, M, H) accounted for by changing the GlobalSamplingFraction parameter (0.3, 0.22, 0.18, respectively) (see Supplemental Information for justification). To account for potential oversampling of Nearctic species, we performed a series of runs where we lowered the number of Nearctic species by randomly pruning 21 (of total 48) Nearctic tips from the tree ten times and performed a BAMM run on each replicate, using the M assumption for the GlobalSamplingFraction parameter.

Using the posteriors generated from these MCMC runs, we sought to 1) explore the overall pattern of *Pheidole* diversification, 2) assess whether there is evidence of diversity regulation, particularly decelerating diversification over time and after colonization of new areas, and 3) test for latitudinal dependency in diversification rate while accounting for phylogenetic non-independence. We visualized the linage specific diversification with the plot.bammdata function from BAMMtools, and the time plot of clade-specific diversification rate was plotted with the plotRateThroughTime function. We used STRAPP (e.g. the traitDependentBAMM function in BAMMtools) to test for significance of any latitude-diversification correlations. We tested for diversification rate vs. either tropicality index or absolute midpoint latitude (one-tailed, 10000 iterations, Spearman’s rho as test statistic). We also checked whether our results were robust to using Pearson correlation as test statistic or coding latitude as a binary variable and using Mann-Whitney test (tropicality>0 or tropicality<0).

#### HiSSE Implementation

The HiSSE approach (Beaulieu & O’Meara, 2016) extends the BiSSE (Binary State Speciation and Extinction model) (Maddison *et al.*, 2007) framework with two advances. First, the HiSSE model itself allows for more complex models in which macroevolutionary rates can be the function of the focal trait and a hidden state. Thus, if our focal character has states 1 and 0 (in our case tropical and extratropical), there could be an influence of a second unobserved character (with states A and B) on a macroevolutionary rates lambda and mu (λ_0A_, λ_0B_, λ_1A_, λ_1B_). Second, importantly, it allows the fitting of null character-independent models (CID) in which a hidden factor(s) drives diversification rate changes without the influence of the focal trait under investigation. This allows trait-dependent BiSSE models to be compared to a character-independent model of similar complexity (CID-2, with two hidden states A and B) and more complicated HiSSE models to be compared to models of similar complexity (CID-4, with four hidden states A, B, C, D). BiSSE (trait dependent speciation-extinction), HiSSE (trait-dependent speciation/extinction with hidden states that also affect speciation/extinction) and CID (trait-independent models with hidden states that affect speciation/extinction) are best used together and models with all structures can be compared.

We fit a range of models with increasing complexity, starting with the BiSSE family of models under the following sets of constraints on the parameters: all diversification and transition rates equal among states, diversification equal but transition rates different (i.e. speciation and or extinction changes with latitude, but transition rates among temperate and tropical are equal), diversification different but transition rates equal (i.e. speciation and extinction vary with state, but transition rates are equal), or all rates free unconstrained to vary with state (the full BiSSE model).

The HiSSE models allow speciation/extinction/transition rates to vary with the focal trait and also among two hidden traits. One question in implementing HiSSE is how to set the transition parameters among states (combination of observed 0/1 and hidden A/B states, with combined state space 0A, 1A, 0B, 1B). We followed suggestions of the authors of the method (Beaulieu & O’Meara, 2016), either setting all transition rates to be equal, or assumed a three parameter rates in which transitions between the observed states could vary but transitions between hidden states is a single parameter.

The CID-2 and CID-4 models are fit including 2 or 4 hidden states, respectively, but with no dependence on the observed traits. The CID-2 is a null model of similar complexity to the full BiSSE model, and the CID-4 is a null model of similar complexity to the HiSSE model. For these, we also assumed alternatively a single rate for all state transitions (observed and hidden) or a three-rate model including two rates for transitions between the observed states and one between all hidden states.

We implemented all of the above analyses using functions in the R package hisse (Beaulieu & O’Meara, 2016). As with the BAMM analysis, we ran all models using either the (L, M, H) assumptions about sampling completeness, and for the global *Pheidole* and New World only. For the New World analyses, we additionally adjusted the sampling fraction (M*) to account for possible undersampling of tropical species relative to extratropical species. As the ML optimization does not always find the global minimum from a single starting point, we ran 20 ML searches for each model using random starting parameters chosen from a uniform distribution on the interval (0,1). For all the models above, we ran them alternatively assuming a fixed root in the tropical state, or root probability estimated with the default “madwitz” method based on the data. As we found the results were insensitive to the root method, we only present results with the default option. After all BiSSE, HiSSE, and CID models were inferred, we compared all models with Akaike’s Information Criterion (AICc) scores.

#### FiSSE Approach

FiSSE (Rabosky & Goldberg, 2017) is a nonparametric test for trait-dependent speciation rates that does not assume an underlying model structure, but rather depends on distributions of branch lengths in the different states. FiSSE is complementary to the BiSSE and is robust to Type-I error. We performed both one-tailed and two-tailed tests of FiSSE to test for speciation differences between temperate and tropical taxa, using the global *Pheidole* and only the New World *Pheidole*. We also performed FiSSE on a set of trees for the New World only where temperate species were thinned to account for possible undersampling of the tropics (see supplemental).

#### Phylogenetic niche conservatism

We performed additional analyses to evaluate the degree to which latitudinal affinity is conserved in *Pheidole*. For this, we first calculated two measures of phylogenetic signal—Blomberg’s K (Blomberg *et al.*, 2003) and Pagel’s lambda (Pagel, 1999)—treating absolute latitudinal midpoint as a continuous trait, using the phylosig() function in the R package *phytools* (Revell, 2012). Second, to estimate the overall evolutionary rates, we fit models of discrete character evolution (treating latitudinal affinity as a binary variable) using the fitDiscrete() function in the R package *geiger* (Pennell *et al.*, 2014). To visualize the evolution of latitudinal affinity, we performed 100 stochastic character maps on the empirical tree using the make.simmap() function, and plotted a summary of state probabilities with the function densityMap(), both from the *phytools* package. Finally, to estimate whether the inferred rate of evolution combined with tropical ancestral state is consistent with the observed richness difference even in the absence of diversity regulation and diversification rate differences, we simulated niche evolution on the empirical tree and maximum likelihood model with the sim.history() function from *phytools*. While tree shape and trait state are not necessarily independent (i.e. the dependent model is implemented in the HiSSE analysis), this analysis asks whether we would be likely to observe a gradient even if they were independent, given that *Pheidole* likely has a tropical ancestor and given the rate that latitudinal affinity evolves. *Pheidole* likely has a tropical ancestor as the most basal *Pheidole* species, *P. fimbriata*, and the sister lineage of *Pheidole, Cephalotes*+*Procryptocerus*, are tropical (Moreau, 2008; Ward *et al.*, 2015).

## RESULTS

*Pheidole* exhibits a latitudinal diversity gradient that is overall similar to ants as a whole (Fig. 1). The BEAST analysis inferred a phylogeny whose major features are consistent with previous studies (Figs. 2, S1). The crown age of the group (i.e. the mrca of *Pheidole fimbriata* with the rest of *Pheidole*) is inferred here to be younger than in a previous study (~29mya vs. ~37mya in Economo *et al.*, 2015a), although closer to the crown age inferred in other recent broader scale phylogenies (Ward *et al.*, 2015).

**Figure 1:**
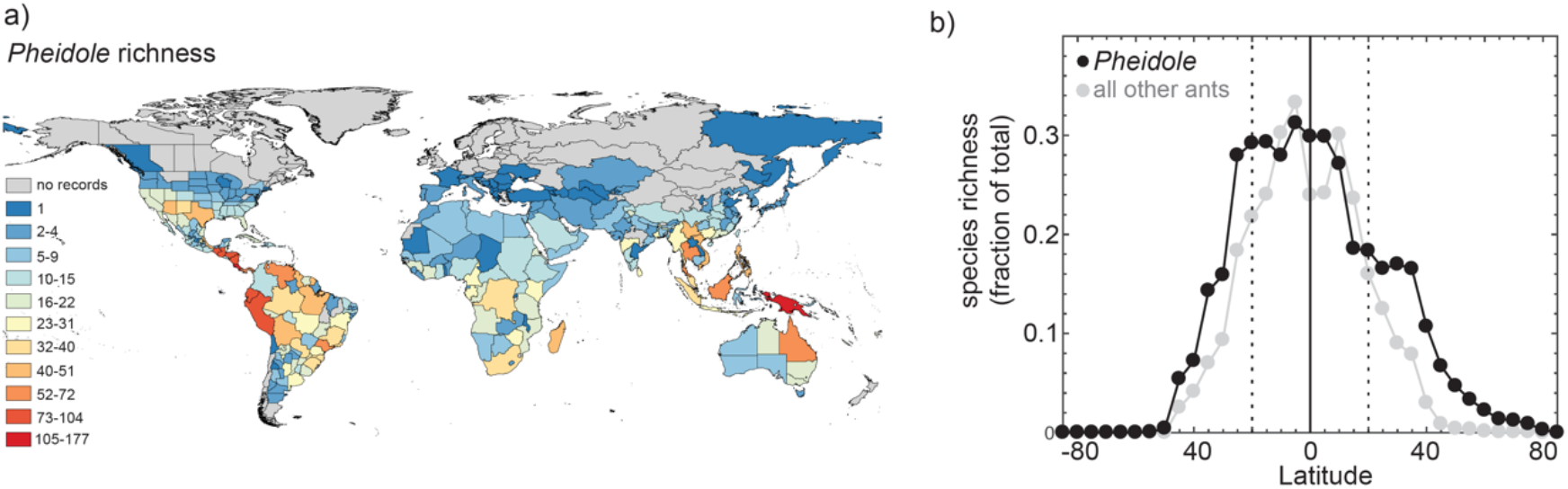
Global patterns of *Pheidole* species richness plotted by a) geographic region and b) 5-degree latitudinal band for 1138 described species/subspecies and 361 morphospec¡es. For comparison, latitudinal distribution of 13771 ant species excluding *Pheidole* are also depicted. Latitudinal richness is expressed as fraction of total richness (either 1499 or 13,771 for *Pheidole* or all other ants, respectively).

According to the BAMM analysis, the hyperdiversification of *Pheidole* began after an acceleration approximately 15-16 mya, and all species except for two basal lineages (*P. fimbriata* and *P. rhea*) are descended from this lineage. Diversification initially occurred in the New World, exhibiting a decelerating trend over time. Around 13mya, a single lineage colonized the Old World and this was associated with another burst of diversification followed by a slowdown in a clade encompassing Asia and Africa. Madagascar and Australia-NG were later colonized, followed by accelerations and subsequent decelerations in each clade (Figs. 2, S1, S2). There were several other accelerations that were not obviously associated with geographic transitions, including one clade in the new world and the *megacephala* group in Afrotropics. This general pattern of sequential colonization-acceleration-deceleration pattern is robust to changing the sampling fraction parameter, although as one would expect the inferred degree of deceleration becomes less pronounced if one assumes that more species are left to be sampled.

**Figure 2:**
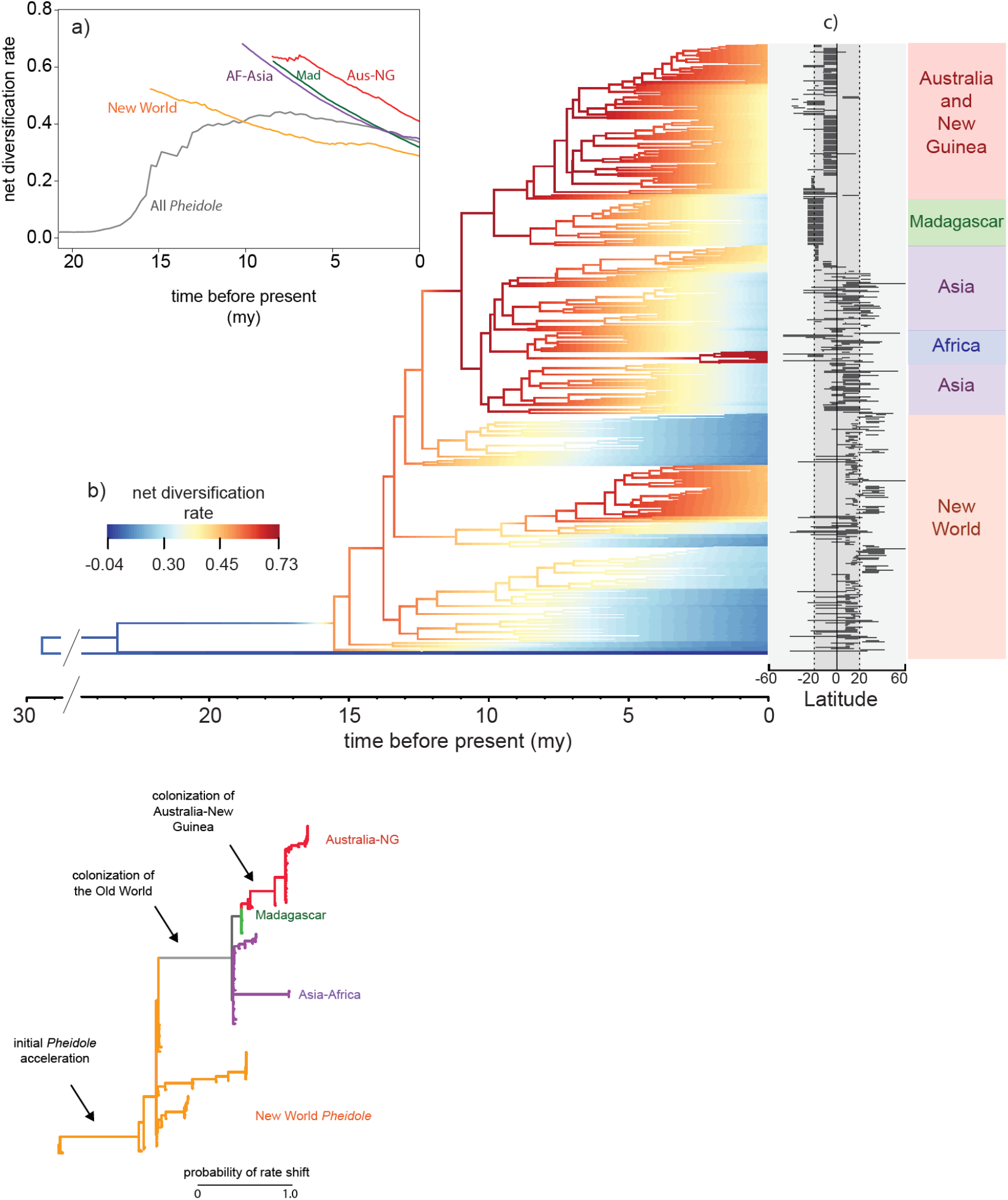
Diversification rate dynamics inferred with BAMM from a phylogeny of 449 Pheidole species, a) Median diversification rates through time of the major *Pheidole* clades. The New World median excludes the two basal species (*P. rhea* and *P. fimbriata*) that fall outside the initial acceleration of *Pheidole* diversification. b) The mcc phylogeny colored with inferred net diversification rate, c) Latitudinal extent of all 449 taxa included in the tree. A high-resolution version with taxon names visible is presented in Figure S1. d) Probable locations of diversification rate shifts. Here, branch length is proportional to probability of a shift.

The extratropical lineages generally belong to young clades nested within larger tropical clades (Figs. 2, S1). While diversification rate varies across the genus to a degree, we could not detect a significant correlation (assessed with STRAPP) between BAMM-inferred net diversification rate and either absolute midpoint latitude or tropicality index across any of the analyses we performed (Fig. 3). These results were similar across variation in the assumed global sampling fractions, whether we calculated correlations for individual clades or the whole tree, and including trees where Nearctic species were culled to account for possible uneven sampling. Although significance tests were one-tailed for higher diversification in the tropics, we also note that none of the observed correlation coefficients were outside the null range in either direction.

**Figure 3:**
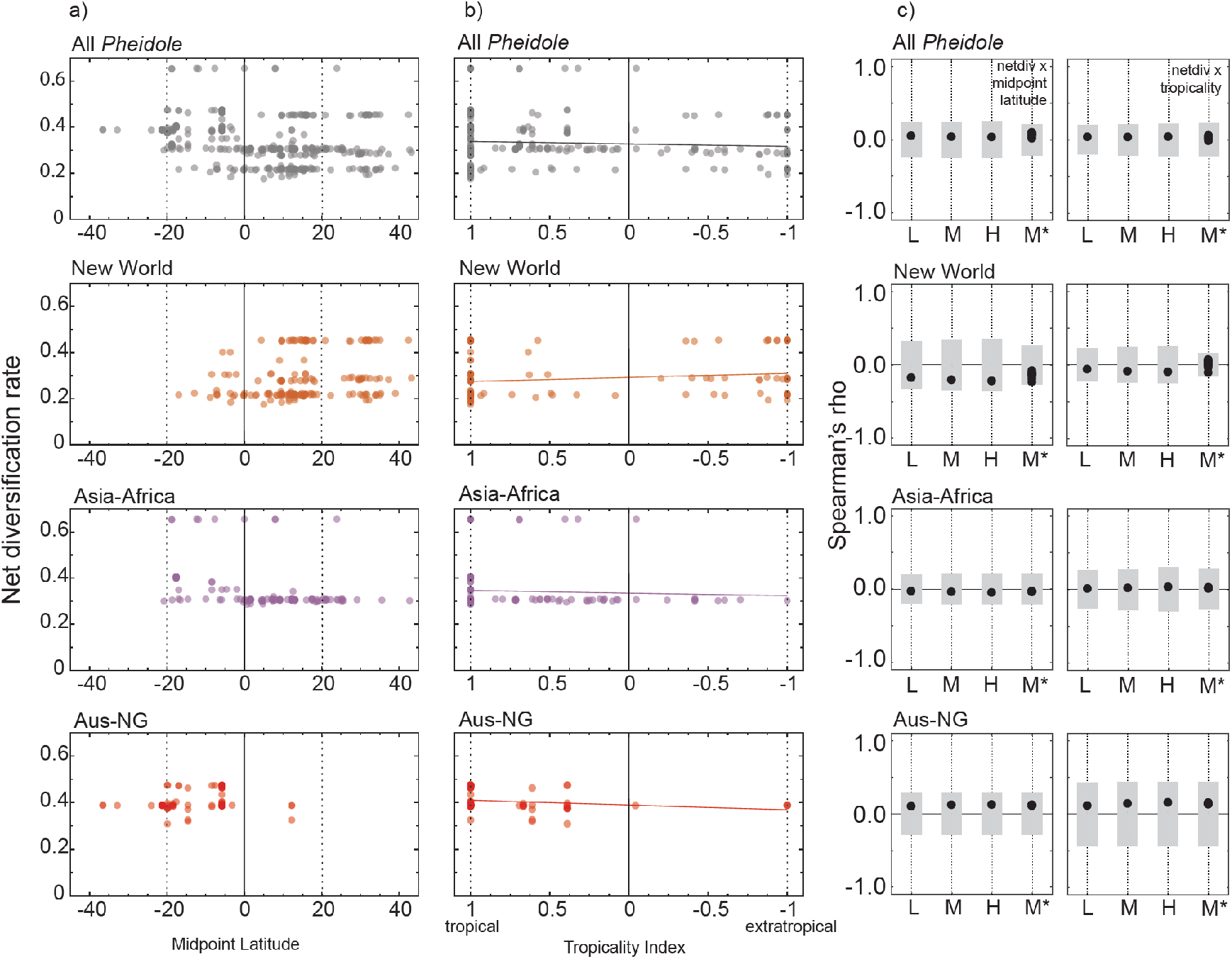
Net diversification rate inferred with BAMM as a function of latitude. Diversification rate of each *Pheidole* species (present day) inferred with BAMM using the “M” assumption of sampling completeness per species a) as a function of latitudinal midpoint and b) tropicality index, which varies from -1 for a species with a range located competely outside the tropics to 1 for a species confined to the tropics, c) Spearman correlations (black dots) for net diversification and either absolute midpoint latitude (left) or tropicality (right), where the grey boxes reflect 95% null distribution generated with STRAPP. L, M, H, reflect different assumptions about unsampled species (low, medium, high estimates of total numbers of *Pheidole*), while M* are 10 trees where temperate species have been culled to account for possible sampling bias (see methods).

The HiSSE analysis was also broadly consistent with BAMM analysis in finding no statistical support for a correlation between macroevolutionary rates and latitude. In general, the CID-2 trait-independent null model outperformed the BiSSE trait-dependent models, and the CID-4 null outperformed the HiSSE trait-dependent models, and the CID-4 models had the global minimum AICc across the different permutations of the analysis (Table 1). Thus, this analysis provided no evidence for latitude-dependent macroevolution in this genus. It is worth noting as well that the AIC-minimizing versions of BiSSE and HiSSE models, which again were themselves not preferred over the null models, generally did not support higher diversification rate in the tropics. The BiSSE model detected a slightly higher diversification rate in the extratropical zone and the HiSSE model either fit models where tropical diversification was higher than extratropical while in one hidden state and lower in the other hidden state, or where the extratropical diversification was always higher in both hidden states. For the New World, use of the sampling effort correction removed this slight, and non-significant difference. The GeoSSE analysis, presented in the supplement, showed overall similar results to BiSSE, a positive latitude-diversification rate trend in the New World, but not global, *Pheidole* that is not robust to the correction for latitudinal undersampling, with some differences in the dispersal pattern inferred probably due to differences the way geographic states are coded. We can only assume that if a CID-like null model were available for GeoSSE, it would also perform better than GeoSSE as it did for BiSSE.

**Table 1:** Summary of delta AICc from the BiSSE and HiSSE trait-dependent models, and the two null models, CID-2 and CID-4. CID-2 is similar in model complexity to the BiSSE model, while CID-4 is similar in model complexity to the HiSSE model. The models were run with different parameter constraints listed below. The L, M, H, refer to the low, medium, and high estimates of missing taxa. M* includes a correction for possible oversampling with latitude. The AICc minimizing model for each analysis is highlighted in bold.

|  |  |  |  |  |  |  |  |  |
| --- | --- | --- | --- | --- | --- | --- | --- | --- |
| Model | Description/constraint | Global *Pheidole*<br>($\Delta AICc$) | | | New World *Pheidole*<br>($\Delta AICc$) | | | |
|  |  | L | M | H | L | M | H | M* |
| BiSSE | Div. rates and transition rates equal across latitudes | 69.9 | 67.4 | 69.6 | 18.3 | 23.2 | 27.8 | 23.2 |
| BiSSE | Div. rates equal, transition rates vary with latitude | 69.4 | 66.9 | 69.2 | 19.4 | 24.2 | 28.9 | 22.3 |
| BiSSE | Div rates vary, transition rates equal with latitude | 69.9 | 67.4 | 69.6 | 18.3 | 23.2 | 27.8 | 23.2 |
| BiSSE | Div. rates and transition rates vary with latitude (full BiSSE model) | 73.5 | 71.1 | 73.3 | 11.6 | 16.1 | 18.9 | 23.2 |
| CID-2 null | 2 hidden states, 1 transition parameter | 33.9 | 21.9 | 17.3 | 0.5 | 2.8 | 5.7 | 2.9 |
| CID-2 null | 2 hidden states, 3 transition parameters | 21.2 | 20.1 | 19.4 | 2.4 | 5.3 | 8.6 | 4.5 |
| HiSSE | Div rates vary with latitude and two hidden states, 1 transition parameter | 36.1 | 21.1 | 16.5 | 0.9 | 0.9 | 0.3 | 8.6 |
| HiSSE | Div rates vary with latitude and two hidden states, 3 transition parameters | 22.5 | 28.0 | 27.7 | 0.2 | 2.0 | 2.3 | 9.5 |
| CID-4 null | Div rates vary with four hidden states, 1 transition parameter | 22.5 | 11.1 | 4.6 | 1.6 | **0.0** | **0.0** | **0.0** |
| CID-4 null | Div. rates vary with four hidden states, 3 transition parameters | **0.0** | **0.0** | **0.0** | **0.0** | 1.1 | 0.4 | 5.7 |

The FiSSE analysis was also consistent with the other analyses in showing no correlation between speciation rate and latitude for the global *Pheidole* (λ_temp_=0.28, λ_trop_=0.27, two tailed p>0.88), but a positive speciation-latitude correlation for the New World alone (λ_temp_ =0.30, λ_trop_ =0.20, two-tailed p<0.026). However, when we dropped extratropical tips from the phylogeny to simulate potential latitudinal undersampling of the tropics, this difference was much more modest and no longer significant (n=10, mean λ_temp_=0.24, S.E.=0.005, mean λ_trop_=0.20, S.E. = 0.0002, p range: 0.19-0.72 among replicates).

The extratropical lineages are clustered with each other on the tree, although it is clear there were numerous transitions out of the tropics (Fig. 4). The tests for phylogenetic signal in latitudinal affinity for Blomberg’s K (K=0.34, p<0.002) and Pagel’s lambda (*λ*=0.95, p<10^−57^) were both highly significant. Symmetric and asymmetric models of discrete character evolution both fit the data comparably well (symmetric model *q_trop->etrop_=q_etrop->trop_*=0.015, AICc=235.5, asymmetric model *q_trop->etrop_*= 0.013, *q_etrop->trop_*=0.060, AICc=234.9). Simulations of character evolution on the empirical phylogeny show that a latitudinal gradient is the most common outcome if one assumes a tropical ancestor and either model for the inferred rate of evolution of latitudinal affinity (Fig. 4).

**Figure 4:**
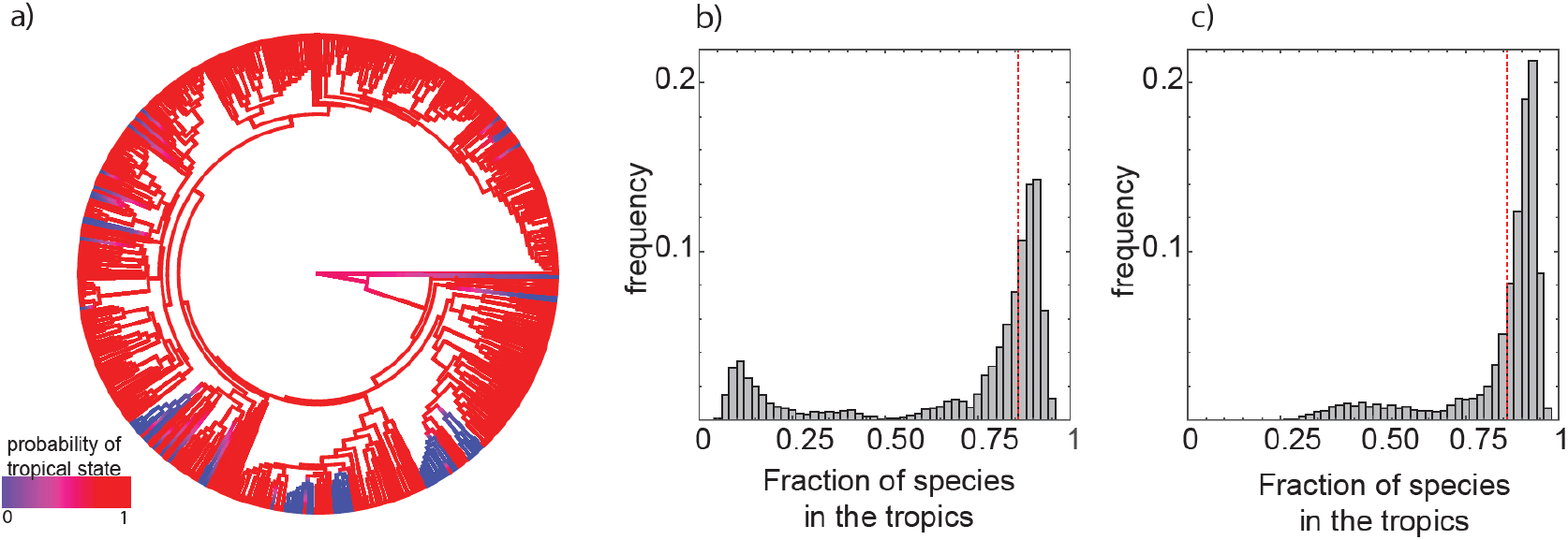
Evolution of latitudinal affinity in *Pheidole*. a) Branch-wise probability of ancestral tropical state inferred from stochastic character mapping, b-c) Histograms of latitudinal richness differences beween tropics and extratropics simulated with stochastic character mapping on the empirical phylogeny assuming a tropical ancestor and the inferred degree of niche conservation using symmetric (b) or asymmetric (c) models of character evolution. The vertical dashed line is the empirical richness fraction.

## DISCUSSION

Our analysis of *Pheidole* macroevolution sheds light on the mechanisms responsible for the evolution of the latitudinal diversity gradient in ants. By focusing on the recent evolutionary dynamics of a single large radiation, our study complements taxon-wide studies that focus on differences among highly divergent clades and deeper timescales (e.g. Cardillo *et al.*, 2005; Weir & Schluter, 2007; Jetz *et al.*, 2012; Pyron & Wiens, 2013; Rolland *et al.*,2014; Kerkhoff *et al*. 2014, Duchêne & Cardillo, 2015; Economo *et al*. In Press).

We find no evidence of higher diversification rate for tropical *Pheidole* lineages across any of our analyses (Figs. 2–4, S1), as would be predicted by the Diversification Rate Hypothesis. In general, the signal of latitude as a trait affecting macroevolutionary rates in the BAMM, HiSSE, and FiSSE analyses was weak to non-existent. When there was some hint of a correlation, for example in the best fitting (but still not better than null) HiSSE/BiSSE analyses, and the FiSSE analysis for New World speciation rate uncorrected for latitudinal sampling bias, it was in the direction of higher diversification/speciation in the temperate zone. However, those correlations were not robust to reasonable assumptions about undersampling in the tropics, thus the overall picture is a lack of evidence for latitudinal dependency for macroevolutionary rates.

We do not view our analysis as ruling out that such systematic macroevolution-latitude relationships may exist, even in *Pheidole*. Rather, our analysis only suggests that such relationships are not the causal factor in the gradient. The Diversification Rate Hypothesis is predicated on the fact that lineages reach different latitudes early on in their evolution, but the disparity of richness is due to different accumulation rates over time. If niche conservatism is too high for lineages to evolve out of the tropics (or vice versa) early on in the radiation, there may be no chance for any latitude-macroevolutionary rate correlations to manifest and be statistically detectible. Thus, we view our analysis as stronger evidence that a diversification rate-latitude correlation is not causal in the latitudinal gradient in *Pheidole*, rather than showing that no such relationship exists in *Pheidole* which could affect biodiversity patterns under the right conditions.

While there is no evidence for relationship between diversification rate and latitude, the pattern of diversification suggests *Pheidole* evolution is being shaped by diversity regulation. Even if one assumes that there are over 1300 undescribed *Pheidole* species (the higher end of our estimated range) in addition to the 1175 currently described species, our analysis found that diversification is still decelerating in the genus. Moreover, there is evidence that diversification accelerated after colonization of new areas, specifically when *Pheidole* colonized the Old World, and again after it colonized Australia and New Guinea. This lends further support to the idea that there are ecological limits to *Pheidole* diversity, because when new continents are colonized, ecological opportunity is high. However, it is more difficult to determine if these limits vary with latitude in a way that is causally responsible for the richness gradient. For example, the limits to diversity could be similar at all latitudes, but phylogenetic conservatism could be causing the higher latitudes to lag behind tropical latitudes in reaching their steady state. Here we might expect a positive trend of *Pheidole* diversification rate running counter to the richness gradient, because temperate zones would be further from the equilibrium number of species. While there were some hints of a positive latitude-diversification correlation (e.g. Supplemental Fig. S2), there was not a robust and statistically supported relationship. A future direction would be to examine how niche overlap and coexistence in *Pheidole* varies with latitude or energetic constraints, as has been pursued in better studied taxa such as birds (Pigot *et al.*, 2016).

Overall, the results match the predictions of the tropical conservatism hypothesis (TCH). We found that latitudinal affinity is moderately conserved in *Pheidole*. While there have been a number of transitions from the tropics to the temperate zone, latitudinal affinity evolves slowly enough to make a richness gradient the most likely outcome simply due its tropical ancestry and phylogenetic inertia. Thus, our study joints a series of recent studies supporting the TCH for woody plants (Kerkhoff *et al.*, 2014), birds (Duchêne & Cardillo, 2015), mammals (Buckley *et al.*, 2010), and butterflies (Hawkins & DeVries, 2009).

These results for *Pheidole* evolution over the last 30my connect well to results on ant diversification on deeper timescales (Economo, In Press), and together tell a coherent story about the evolution of latitudinal gradients in ants. Most ant lineages older than 34mya are reconstructed to be tropical, including the *Pheidole* stem lineage. Around 15 mya, *Pheidole* exhibited a many-fold acceleration in diversification rate and began a massive radiation. The reason for this initial acceleration, such as evolution of a key innovation, remains unknown. It took time for some *Pheidole* lineages to evolve the requisite traits for colonization of high latitudes. Once colonization of cold biomes occurred, diversification was not detectibly slower. In their analysis across all ant clades, Economo et al. (In Press) also found no evidence for elevated net diversification rates among clades centered in the tropics relative to those in the temperate zone, although clades are quite heterogeneous in rate, probably due to other latent biological and historical differences. It remained possible that diversification rate is correlated with latitude within the large clades, but biological differences among clades obscured this pattern. Within *Pheidole*, diversification rate is much less heterogeneous, but there is still no evidence of a negative latitudinal correlation, implying that lack of phylogenetic resolution within large clades was not hiding this relationship in the previous analysis (Economo et al. *In Press). Pheidole* provides additional insight that diversity regulation is a prominent feature of the global evolution of the genus, although it is unclear if it is a causal factor in the gradient. Finally, since we have a better handle on sampling biases within *Pheidole* than we do for ants as a whole, we can be more confident that latitudinal sampling biases are not masking latitudinal diversification rate variation.

While thus far the evidence is consistent with both phylogenetic niche conservatism (TCH) and diversity regulation (ERH) playing a role in *Pheidole* diversification, determining whether one alone or both together are responsible for the diversity gradient remains a challenge for future work. Moreover, as these are “umbrella” hypotheses, each individual hypothesis could encompass a range of different mechanisms. One way forward is a hierarchical, systematic approach, where broad categories of hypotheses are evaluated (e.g. like these in this study), followed then by more targeted studies devised to tease apart the mechanisms within the larger classes of hypothesis that fit the data well. We also agree with the approach advocated by Hurlbert and Stegen (Hurlbert & Stegen, 2014a), toward a quantitative formulation of multiple competing and intersecting hypotheses, combined with a simulation-based approach to identify their key predictions. We felt initial efforts in this direction were not yet mature enough to use as a basis for the current study, but look forward to further development of the approach in the future. Finally, we need further work to resolve and analyze other hyperdiverse ant radiations (e.g. *Camponotus, Strumigenys, Tetramorium*) that also exhibit strong latitudinal gradients.

Despite the high level of research effort directed toward understanding the latitudinal gradient, the matter is far from resolved (Mittelbach *et al.*, 2007). Studies have differed in their conclusions about the origins of the gradient, probably due to both differences in conceptual and methodological approaches and real variation in process and history across taxonomic groups. The former should continue to improve as we develop more penetrating quantitative methods that make use of more diverse data types. Variability across taxonomic groups is best assessed and understood by examining more of them. The vast majority of studies on the diversity gradient have focused on vertebrates. While of obvious intrinsic interest, vertebrates may not be good surrogates for understanding general patterns across the rest of the tree of life. For example, mammals have been impacted by human activities so dramatically that it can affect large-scale macroecological patterns (Turvey & Fritz, 2011; Santini, 2017). With development of global invertebrate datasets like the one analyzed here, we stand to broaden our perspective on large-scale biological patterns and their origins.

## DATA ACCESSIBILITY

Molecular sequences have been deposited to GenBank (see Table S1 for accession numbers). We have also provided the alignment, BEAST xml file, and geographic dataset in a supplemental data archive. The GABI dataset can be accessed on the interactive website antmaps.org.

## ACKNOWLEDGEMENTS

This work was supported by NSF (DEB-1145989 to EPE and LLK), by subsidy funding to OIST, and by a Japan Society for the Promotion of Science Kakenhi grant to EPE. We thank P.S. Ward and B.L. Fisher for sharing specimens and for all the data contributors to the GABI project.

## BIOSKETCH

The research team is interested in the ecology and evolution of biodiversity, especially insects.

